# Discovery of a CI-994 derivative as a dual modulator of class I HDACs and Wnt/β-catenin signaling for Alzheimer’s disease therapy

**DOI:** 10.64898/2026.04.30.721954

**Authors:** Wenyan Lu, Thomas R. Caulfield, Eunmi Lee, Suren Jeevaratnam, Ni Wang, Guojun Bu, Takahisa Kanekiyo, Yonghe Li

## Abstract

Alzheimer’s disease (AD) is a multifactorial disease with mixed pathologies. Consequentially, drugs targeting multiple pathological processes may offer synergistic benefits. While histone deacetylase (HDAC) inhibitors have demonstrated efficacy in alleviating AD-related pathologies in animal models, the neuroprotective Wnt/β-catenin signaling pathway remains compromised in AD brain. CI-994 is a class I HDAC inhibitor containing N-(2-aminophenyl)-benzamide. Our recent studies indicate that CI-994 is also an activator of Wnt/β-catenin signaling by stabilizing Wnt co-receptor LRP6. We herein use CI-994 as a scaffold to develop novel potent dual modulators of class I HDACs and Wnt/β-catenin signaling for AD therapy. Our lead compound, W2A-28, selectively inhibits class I HDAC1, 2 and 3 with IC_50_ values of 0.51 μM, 0.68 μM, and 0.22 μM, respectively, and shows no inhibitory activities on other HDACs. Furthermore, W2A-28 potently activates Wnt reporter activity with an EC_50_ value of 1.61 μM in Wnt-3A-expressing HEK293 cells. As expected, activation of Wnt/β-catenin signaling by W2A-28 is associated with elevated LRP6 protein level. Importantly, W2A-28 displays excellent microsomal stability in both mouse and human liver microsomal stability assays, alongside high permeability and a lack of active efflux in MDR1-MDCKII models. Critically, W2A-28 treatment significantly enhances histone acetylation, activates Wnt/β-catenin signaling, and suppresses tau phosphorylation in AD patient-specific cerebral organoids carrying *APOE* ε4/ε4 or *APOE* ε3/ε4 with *PSEN1* M146V mutation. Our findings position W2A-28 as a promising multi-target drug candidate for AD therapy.

## Introduction

Alzheimer’s disease (AD) is a complex neurodegenerative disorder associated with multiple pathological mechanisms, and multi-targeted therapies are desirable for this incurable disease (1, 2). Histone deacetylases (HDACs) are essentia epigenetic regulators that directly regulate synaptic plasticity, learning, and memory formation in the brain (3-7), and small molecule inhibitors of class I HDACs can suppress AD pathologies and alleviate memory deficits in mouse AD models (8-14). Wnt/β-catenin signaling, which is greatly suppressed through multiple dysregulation mechanisms in AD brain, is crucial for synaptic plasticity, neurogenesis and blood-brain barrier integrity and function (15). Mounting evidence supports that restoration of Wnt/β-catenin signaling is a promising therapeutic strategy for AD (15).

CI-994 (Tacedinaline) is a specific class I HDAC inhibitor and has undergone multiple phase II/III clinical trials for advanced nonsmall cell lung cancer, pancreatic cancer and multiple myeloma (16-18). CI-994 is oral bioavailable and brain penetrable (19, 20), and can function as a molecular memory aid and alleviate cognitive impairment in several mouse models of neurodegenerative disorders (8, 20-24). CI-994 improves cognitive function in mouse models of aging and Birk-Barel intellectual disability syndrome (22, 23), and reduces neuronal loss and enhances functional recovery from spinal cord injury in mice (21). Importantly, CI-994 reduces tau pathology in AD patient-specific induced pluripotent stem cell (iPSC)-derived cerebral organoids (14) and alleviates amyloid pathology in APP/PS1 mice (8).

We have recently reported that CI-994 acts as a dual modulator of class I HDACs and Wnt/β-catenin signaling and displays profound effects on suppressing tau phosphorylation and modulating the expression of genes associated with synapse plasticity and cognitive function in AD patient-specific iPSC-derived neurons and cerebral organoids (14). In the present study, we use CI-994 as a scaffold to develop novel dual modulators that inhibit class I HDACs and activate Wnt/β-catenin signaling activation and identify W2A-28 as a leading candidate for the treatment of AD.

## Materials and Methods

### Compounds

CI-994 (cat. no. S2818) and niclosamide (cat. no. S3030) were purchased from Selleck Chemicals LLC. W2A-28, W2A-42 and W2A-43 were designed at Mayo Clinic Jacksonville and synthesized by Enamine US Inc (Monmouth Jct., NJ 08852) with the purity >95%.

W2A-28: 1H NMR (DMSOd6): δ = 10.50 (s, 1H, *NH*), 9.94 (s, 1H, *NH*), 8.86 (m, 1H, *C6pyrH*), 8.25 (m, 1H, *C4pyrH*), 8.09 (d, 1H, *C3pyrH*), 7.50 (d, 1H, *CArH*), 6.95 (t, 1H, *CArH*), 6.82 (d, 1H, *CArH*), 6.65 (t, 1H, *CArH*), 4.96 (s, 2H, *NH2*), 2.13 (s, 3H, *CH3*) ppm. LS-MS (m/z): calcd./found for [M+H]+ 271.12/271.2.

W2A-42: ^1^H NMR (DMSOd6): δ = 10.86 (s, 1H, *NH*), 9.18 (s, 1H, *C*_*6pyr*_*H*), 9.06 (q, 1H, *NH*), 8.47 (d, 1H, *C*_*4pyr*_*H)*, 8.23 (d, 1H, *C*_*3pyr*_*H)*, 7.65 (d, 1H, *C*_*Ar*_*H*), 7.57 (d, 1H, *C*_*Ar*_*H*), 7.45 (t, 1H, *C*_*Ar*_*H*), 7.37 (t, 1H, *C*_*Ar*_*H*), 5.52 (s, 2H, *NH*_*2*_), 2.84 (s, 3H, *NCH*_*3*_) ppm. LS-MS (m/z): calcd./found for [M+H]^+^ 271.12/271.1.

W2A-43: ^1^H NMR (DMSOd6): δ = 10.63 (s, 1H, *NH*), 8.68 (q, 1H, *NH*), 8.17 (d, 2H, *C*_*Ar*_*H)*, 7.99 (d, 2H, *C*_*Ar*_*H)*, 7.57 (d, 1H, *C*_*Ar*_*H*), 7.45 (m, 1H, *C*_*Ar*_*H*), 7.39 (m, 1H, *C*_*Ar*_*H*), 2.75 (s, 3H, *NCH*_*3*_) ppm. LS-MS (m/z): calcd./found for [M+H]^+^ 270.12/270.2.

### Cell culture

HEK293 cells (CRL-1573) and N2a neuroblastoma cells (CCL-131) were obtained from ATCC. Human fibrosarcoma cancer HT1080 cells stably transfected with HA-tagged LRP6 have been described before (25). HEK293 cells were cultured in DMEM medium (Gibco) containing 10% of fetal bovine serum (FBS) (Gemini Bio-Products), 2 mM of L-glutamine, 100 units/ml of penicillin and 100 μg/ml of streptomycin, and N2a cells were cultured in DMEM/Opti-MEM medium (1:1) (Gibco) supplemented with 5% FBS, 2 mM of L-glutamine, 100 units/ml of penicillin and 100 μg/ml of streptomycin. All the cells were grown under standard cell culture conditions at 37 °C in a humidified atmosphere with 5% CO_2_.

### HDAC inhibition profiling and IC_50_ determination

Profiling HDAC inhibition and determination of IC_50_ values of HDAC1, 2 and 3 were performed by Reaction Biology Corp., Malvern, PA 19355. CI-994, W2A-28, W2A-42 and W2A-43 were tested in single dose duplicate at 10 μM against 10 HDACs (HDAC1-9, HDAC11) to obtain HDAC inhibition profile. W2A-28, W2A-42 and W2A-43 were tested against HDAC1, HDAC2 and HDAC3 in 10-dose IC_50_ mode, in singlet, with 3-fold serial dilution starting at 100 μM to determine IC_50_ values by W2A-28, W2A-42 and W2A-43.

### Wnt reporter activity assay

The Super8XTOPFlash luciferase assay was conducted in HEK293 cells transiently transfected with the Super8XTOPFlash luciferase construct, pcDNA3-Wnt3-HA and β-galactosidase-expressing vector as described in our previous studies (26, 27).

### Determination of metabolic stability

Metabolic stability of CI-994, W2A-28, W2A-42 and W2A-43 in mouse and human liver microsomes was measured by Enamine US Inc (Monmouth Jct., NJ 08852). Male BALB/c pooled liver microsomes (XenoTech, cat. no. M3000) and mixed gender human pooled liver microsomes (XenoTech, cat. no. H0630) were used. Propranolol hydrochloride (Sigma-Aldrich, cat. no. P0884), imipramine hydrochloride (Sigma-Aldrich, cat. no. 17379) and diclofenac (Enamine, cat. no. EN300-119509) were served as reference compounds. Metabolic stability of test compounds CI-994, W2A-28, W2A-42 and W2A-43 and reference compounds in mouse and human liver microsomes was analyzed at five time points over 40 minutes using HPLC-MS. Metabolic stability was defined as the percentage of parent compound lost over time in the presence of a metabolically active test system, and intrinsic clearance (Cl_int_) was determined (28).

### Determination of MDR1-MDCKII permeability

MDR1-MDCKII permeability of CI-994, W2A-28, W2A-42 and W2A-43 was measured by Enamine US Inc (Monmouth Jct., NJ 08852). Canine MDR1 Knockout, human MDR1 Knockin MDCKII cells (MDR1-MDCKII) (Sigma-Aldrich, cat. no. MTOX1303) were used. The integrity of the cell monolayers was determined by measuring the trans-epithelial electrical resistance (TEER, Ω × cm2) using an epithelial voltammeter, and the rates of compounds transport in apical (A)-to-basolateral (B) direction (A-B) and in the basolateral (B) to apical (A) direction (B-A) were determined. The effect of the inhibition of P-gp-mediated transport of the tested compounds was assessed by determining the bidirectional transport in the presence or absence of cyclosporine A. The apparent permeability (P_app_) was calculated and expressed in 10^-6^ cm/sec. Efflux ratio of Papp(B-A)/Papp(A-B) was calculated to reveal the difference in P_app_ as a result of active transport. Ketoprofen (Enamine, cat. no. EN300-120644), atenolol (Sigma-Aldrich, cat. no. 74827), digoxin (FLUKA, cat. no. 04599), and quinidine (Sigma-Aldrich, cat. no. Q3625) were used as reference compounds (29-31).

### Generation of iPSCs from human skin fibroblasts, trilineage differentiation of human iPSCs and culture of iPSC-derived cerebral organoids

Human skin biopsies from a female patient with FAD (MC0127) carrying *PSEN1* M146V mutation was obtained from Mayo Clinic patients under IRB protocol with patient consent for research, which was approved by the Mayo Clinic Institutional Review Board. The generation of iPSCs from human skin fibroblasts and trilineage differentiation of iPSCs were performed as described before (14, 32). The *APOE* genotype of iPSC line MC0127 was determined by the TaqMan genotyping assay with probes rs7412 and rs429358 from Thermo Fisher Scientific. The generation of AD patient-specific iPSC line MC0020 carrying *APOE* ε4/ε4 genotype has described before (32). The culture of iPSC-derived cerebral organoids was performed as described in our previous studies (32, 33).

### Cerebral organoid drug treatment and protein extraction

Cerebral organoids at 3 months of age were treated with W2A-28 at 0, 1 and 2 μM for 3 days. Three organoids were pooled as one sample, and a total of 9 samples (3 samples per group) were studied. After treatment, the cerebral organoids were harvested and lysed with RIPA Lysis Buffer (EMD Millipore, cat. no. 20-188) supplemented with Protease Inhibitor Cocktail (Roche, cat. no. 11836170001) and Phosphatase Inhibitor Cocktail (Roche, cat. no. 04906837001). The lysates were sonicated and incubated on ice for 60 min and centrifuged at 21,000 g for 45 min at 4 °C. Supernatants were collected, and total protein concentration was measured using a Bio-Rad Protein Assay Kit (Bio-Rad, cat. no. 5000002EDU).

### Western blotting

Equal amounts of protein of RIPA lysates from HEK293 cells, SH-HA-tagged LRP6-expressing HT1080 cells, N2a cells and cerebral organoids were loaded per well and subjected to SDS-PAGE under reducing conditions, and Western blot analyses were conducted as described before (34, 35). The primary antibodies and their dilutions used in this study are as follows: acetyl-histone H3 (Cell Signaling Technology, 9649S, 1:1000), histone H3 (Cell Signaling Technology, 4499S, 1:1000), β-catenin (BD Biosciences, 610154, 1:5000), NeuroD1 (Cell Signaling Technology, 62953S, 1:1000), phospho-tau (AT8) (Fisher Healthcare, clone AT8, ENMN1020, 1:500), phospho-tau (AT180) (Fisher Healthcare, clone AT180, ENMN1040, 1:500), tau (Fisher Healthcare, clone HT7, clone HT7, ENMN1000, 1:1000), β3-tubulin (Cell Signaling Technology, 5568S, 1:5000), β-actin (Cell Signaling Technology, 3700S and 4970S, 1:5000), and α-tubulin (Cell Signaling Technology, 2125S and 3873S, 1:5000).

### Statistical analysis

All data were presented as mean values ± SEM. Statistical analyses were performed with the GraphPad Prism 9 software. Two-tailed, unpaired t test was used for comparison of two groups, and one-way ANOVA with Dunnett’s/Tukey’s multiple comparison test.

## Results

### W2A-28 displays a similar activity in inhibiting class I HDAcs and is more potent than CI-994 in activating Wnt/β-catenin signaling

We recently reported that CI-994 is a dual modulator of class I HDACs and Wnt/β-catenin signaling (14). CI-994 specifically inhibits class I HDAC1, 2 and 3 activities with IC_50_ values of 0.62 μM, 0.59 μM and 0.47 μM, respectively, and enhances Wnt reporter activity in Wnt-3A-expressing HEK293 cells with an EC_50_ value around 5.82 μM (14). To study the structure-activity relationship (SAR) and optimize CI-994 in modulating HDAC activity and Wnt/β-catenin signaling, we designed, synthesized three CI-994 derivatives W2A-28, W2A-42 and W2A-43 (***Fig. 1A***) and examined their activities. Like CI-994, its derivatives W2A-28, W2A-42 and W2A-43 at 10 μM inhibited HDAC1, 2 and 3 enzyme activity and had no inhibitory effects on other HDACs (***Table 1***). W2A-28 displayed better activity than W2A-42 and W2A-43 in inhibiting class I HDAC1, 2 and 3 with IC_50_ values of 0.51 μM, 0.68 μM and 0.22 μM, respectively (***Fig. 1B***). Importantly, W2A-28 also exhibited better activity than CI-994, W2A-42 and W2A-43 in activating Wnt/β-catenin signaling with an EC_50_ value of 1.61 μM in Wnt-3A-expressing HEK293 cells (***Fig. 1C***). Indeed, W2A-28 at 8 μM was significantly more potent than CI-994, W2A-42 and W2A-43 in upregulating β-catenin levels in neuronal N2a cells (***Fig. 2)***. Together, these findings indicate that W2A-28 is a dual modulator with better activity than CI-994 in Wnt/β-catenin signaling activation.

**Table 1:** Effects of CI-994, W2A-28, W2A-42 and W2A-43 on HDAC enzyme activity.

|  | CI-994 |  |  | W2A-28 |  |  | W2A-42 |  |  | W2A-43 |  |  |
| --- | --- | --- | --- | --- | --- | --- | --- | --- | --- | --- | --- | --- |
|  | Data1 | Data2 | Mean | Data1 | Data2 | Mean | Data1 | Data2 | Mean | Data1 | Data2 | Mean |
| **HDAC1** | 11.23 | 12.47 | **11.85** | 12.00 | 11.65 | **11.82** | 20.65 | 21.24 | **20.95** | 15.53 | 15.68 | **15.61** |
| **HDAC2** | 7.92 | 6.31 | **7.12** | 6.25 | 6.67 | **6.46** | 19.62 | 21.97 | **20.79** | 10.31 | 13.17 | **11.74** |
| **HDAC3** | 2.16 | 2.22 | **2.19** | 3.15 | 4.83 | **3.99** | 7.05 | 6.62 | **6.83** | 5.32 | 4.88 | **5.10** |
| **HDAC4** | 88.96 | 94.73 | **91.85** | 96.38 | 97.70 | **97.04** | 101.09 | 103.14 | **102.12** | 80.17 | 83.27 | **81.27** |
| **HDAC5** | 81.57 | 86.01 | **83.79** | 99.76 | 99.83 | **99.79** | 103.50 | 98.39 | **100.95** | 87.68 | 90.43 | **89.05** |
| **HDAC6** | 82.95 | 81.11 | **82.03** | 89.31 | 92.05 | **90.68** | 93.18 | 99.05 | **96.11** | 87.59 | 89.81 | **88.70** |
| **HDAC7** | 96.83 | 98.12 | **97.47** | 99.39 | 103.13 | **101.26** | 104.57 | 103.11 | **103.84** | 95.88 | 94.66 | **95.27** |
| **HDAC8** | 77.07 | 78.14 | **77.61** | 94.87 | 93.21 | **94.04** | 88.83 | 86.84 | **87.83** | 78.93 | 87.22 | **83.07** |
| **HDAC9** | 79.35 | 82.57 | **80.96** | 93.53 | 99.35 | **96.44** | 84.21 | 88.12 | **86.17** | 85.48 | 81.60 | **83.54** |
| **HDAC10** | ND | ND |  | ND | ND |  | ND | ND |  | ND | ND |  |
| **HDAC11** | 102.96 | 98.70 | **100.83** | 96.49 | 104.21 | **100.35** | 103.21 | 108.66 | **105.93** | 99.35 | 105.48 | **102.42** |

Data are presented as % enzyme activity relative to DMSO controls. CI-994 and niclosamide were tested in duplicate at 10 μM. ND, not determined.

**Fig. 1.**
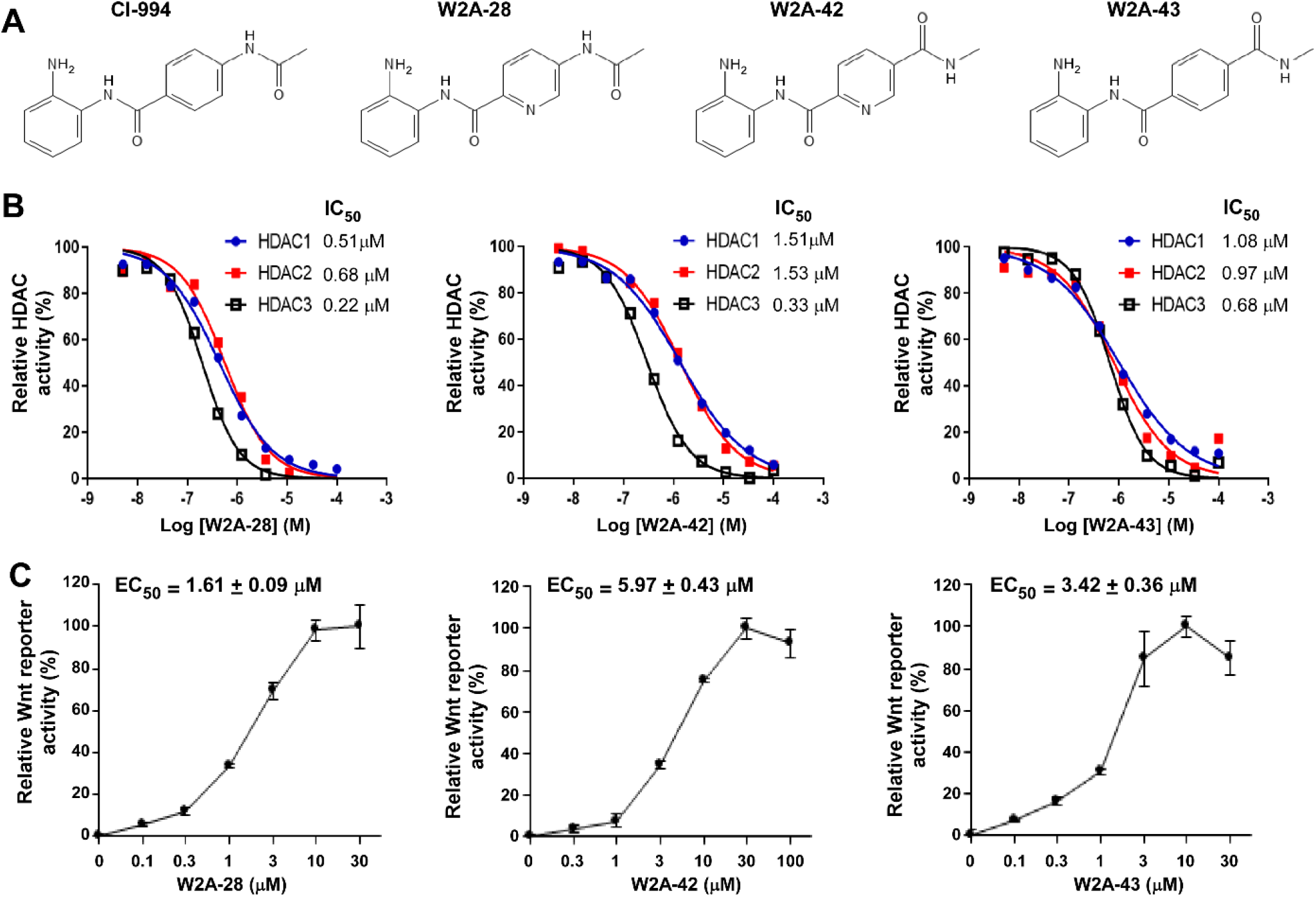
W2A-28, W2A-42 and W2A-43 are dual modulators of class I HDACs and Wnt/β-catenin signaling. (A) The structure of CI-994 and its derivatives W2A-28, W2A-42 and W2A-43. (B) Inhibition of enzyme activity of HDAC1, HDAC2 and HDAC3 by W2A-28, W2A-42 and W2A-43, and IC_50_ values are listed. (C) Activation of Wnt/β-catenin signaling by W2A-28, W2A-42 and W2A-43. Wnt reporter assay in Wnt3A-expressing HEK293 cells with the treatment of W2A-28, W2A-42 and W2A-43 for 24 h. EC_50_ values are shown as mean ± SEM of 3 independent experiments.

**Fig. 2:**
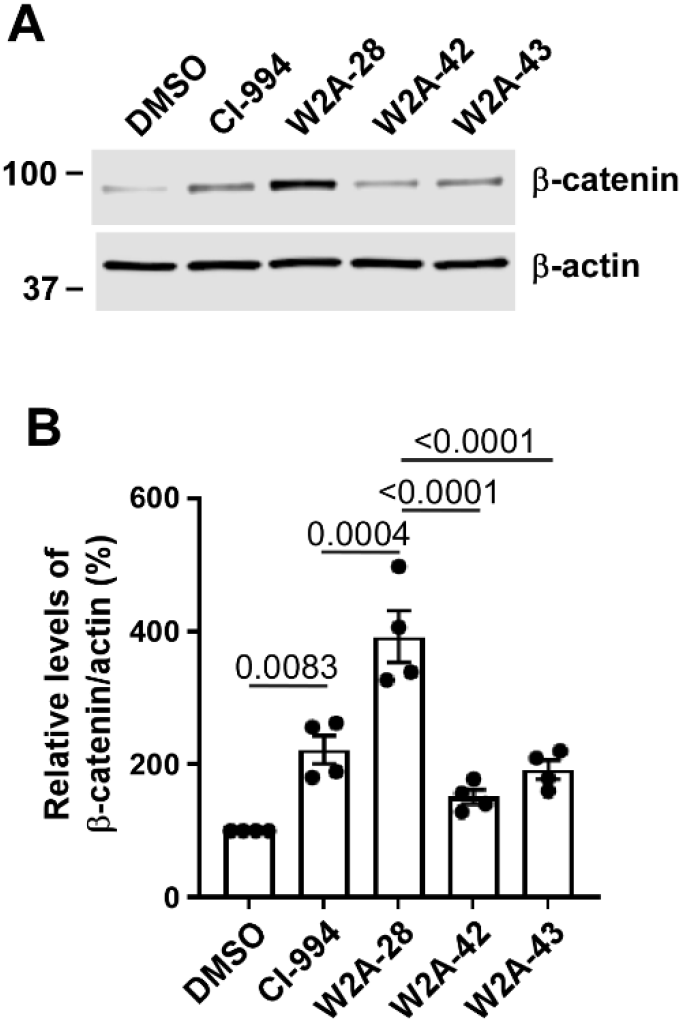
Activation of Wnt/β-catenin signaling by CI-994, W2A-28, W2A-42 and W2A-43 in N2A cells. N2a cells were treated with CI-994, W2A-28, W2A-42 and W2A-43 at 8 μM for 24 h, and the level of β-catenin was examined by Western blotting. Data represent mean ± SEM from four independent experiments. *P* values were calculated using one-way ANOVA with Tukey’s multiple comparison test.

### Activation of Wnt/β-catenin signaling by W2A-28 is associated with upregulation of Wnt co-receptor LRP6

Niclosamide, an analog of CI-994, displays the opposite effect on Wnt/β-catenin signaling by promoting Wnt co-receptor LRP6 degradation (27, 36, 37). It was found that W2A-28 significantly enhanced Wnt co-receptor LRP6 protein level, which is opposite to the effect of niclosamide on LRP6 protein (27, 36, 37), in HA-tagged LRP6-expressing HT1080 cells (***Fig. 3A & 3B***). Moreover, nicotamide at 1 μM abolished W2A-28-induced Wnt reporter activity in Wnt-3A-expressing HEK293 cells (***Fig. 3C***). Together, these findings indicate that upregulation of LRP6 protein is associated with W2A-28-mediated activation of Wnt/β-catenin signaling.

**Fig. 3:**
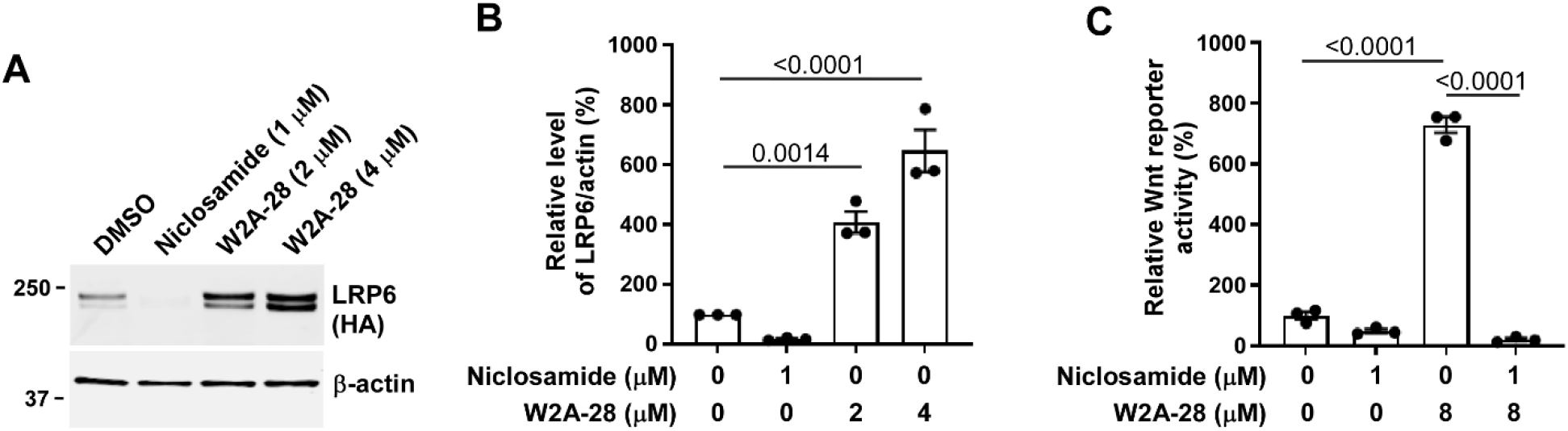
Activation of Wnt/β-catenin signaling by W2A-28 is associated with Wnt co-receptor LRP6 upregulation. (A, B) Western blotting analyses of LRP6 levels in HA-tagged LRP6-expressing HT1080 cells with the treatment of niclosamide (1 μM) and W2A-28 (2 or 4 μM) for 24 h. Data represent mean ± SEM from three independent experiments. (C) Wnt3A-expressing HEK293 cells were treated with niclosamide and W2A-28 at the indicated concentration for 24 h, and Wnt reporter activity was measured. *P* values were calculated using one-way ANOVA with Dunnett’s multiple comparison test (B) or Tukey’s multiple comparison test (C).

### W2A-28 exhibits excellent microsomal stability

Metabolic stability is crucial in the success of drug candidates, and first-pass metabolism is associated with poor oral bioavailability and short half-life (38). Therefore, we examined the metabolic stability of W2A-28 along with CI-994, W2A-42 and W2A-43 in mouse and human liver microsomes. Propranolol and imipramine were served as reference drugs in mouse liver microsomal studies, and propranolol and diclofenac were served as reference drugs in human liver microsomal studies (28). It was found that W2A-28 along with CI-994, W2A-42 and W2A-43 displayed excellent microsomal stability in both mouse and human liver microsomal stability assays (***Fig. 4***), suggesting that W2A-28 could have a good *in vivo* bioavailability.

**Fig. 4.**
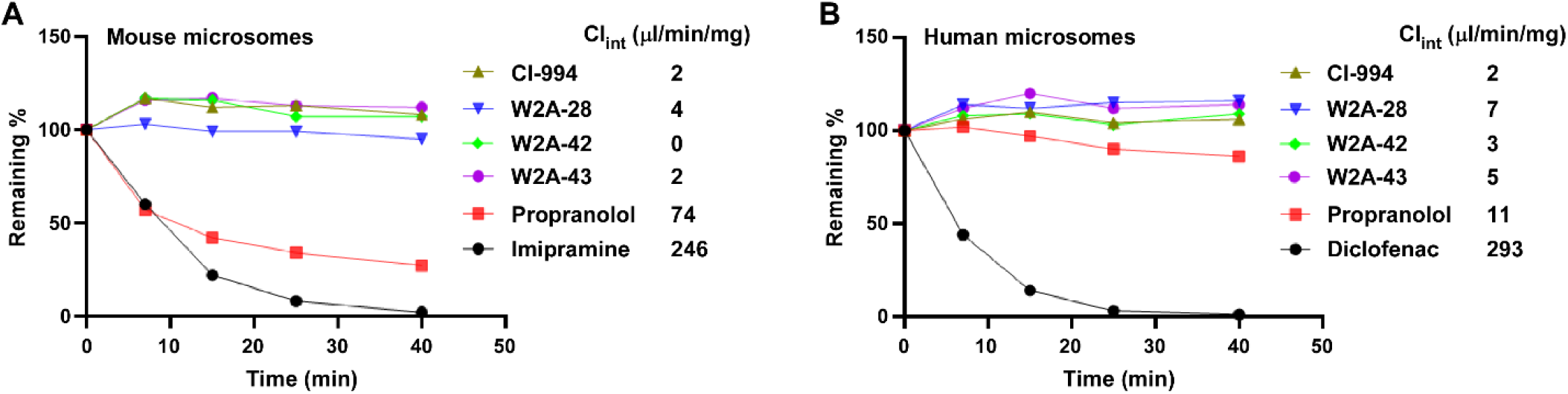
Microsomal stability of CI-994 and its derivatives W2A-28, W2A-42 and W2A-43. The metabolic stability of CI-994, W2A-28, W2A-42, W2A-43 and reference compounds in mouse microsomes (A) and human liver microsomes (B) was analyzed at five time points over 40 minutes. The levels of metabolic stability were measured as the percentage of parent compound lost over time in the presence of a metabolically active test system, and the values of intrinsic clearance (Cl_int_) were calculated and listed.

### W2A-28 displays better permeability than CI-994 in MDR-MDCKII cells

Canine MDR1 Knockout/Human MDR1 Knockin (MDR1-MDCKII) cells are widely used for studying drug efflux, permeability and P-gp substances, and predicting *in vivo* drug brain penetration (39, 40). It was found that the levels of apical-to-basal (A-B) permeability coefficient of W2A-28 and W2A-42 were 25.09×10^−6^ cm/s and 22.01×10^−6^ cm/s, respectively, which were about 2 to 2.5 folds of those of CI-994 and W2A-43 (***Table 2***). In addition, W2A-28, W2A-42 and W2A-43 had an efflux ratio of 0.9 (***Table 2***), indicating that W2A-28, W2A-42 and W2A-43 were not substrates for active efflux transporters. Moreover, cyclosporine A, an inhibitor of P-gp, had no effects on the permeability of all tested compounds in MDR1-MDCKII cells (***Table 3***). Taken together, these findings indicate that W2A-28 along and W2A-42 as well show high permeability in MDR1-MDCKII cells, do not undergo active efflux, and are not a P-gp substrate.

**Table 2:** Apical-to-basal (A-B) and basal-to-apical (B-A) permeability of CI-994, W2A-28, W2A-42 and W2A-43.

| Compound | P<sub>app</sub> (A-B), 10<sup>-6</sup> cm/s |  |  | P<sub>app</sub> (B-A), 10<sup>-6</sup> cm/s |  |  | Efflux ratio* |
| --- | --- | --- | --- | --- | --- | --- | --- |
|  | Data1 | Data2 | Mean | Data1 | Data2 | Mean |  |
| Ketoprofen | 1.30 | 1.34 | **1.32** | ND | ND |  |  |
| Atenolol | 20.71 | 17.50 | **19.10** | 15.24 | 14.89 | **15.06** | **0.79** |
| Quinidine | 5.97 | 4.84 | **5.40** | 30.56 | 27.26 | **28.91** | **5.35** |
| Digoxin | 0.18 | 0.17 | **0.17** | 6.53 | 9.49 | **8.01** | **45.98** |
| CI-994 | 9.40 | 9.09 | **9.24** | 11.20 | 10.71 | **10.96** | **1.19** |
| W2A-28 | 25.90 | 24.27 | **25.09** | 21.68 | 23.59 | **22.63** | **0.90** |
| W2A-42 | 22.23 | 21.79 | **22.01** | 22.23 | 17.92 | **20.08** | **0.91** |
| W2A-43 | 7.38 | 11.50 | **9.44** | 7.88 | 9.26 | **8.57** | **0.91** |

\* Efflux ratio is expressed as the quotient of P<sub>app</sub>(B-A) to P<sub>app</sub>(A-B).

**Table 3:** Apical-to-basal (A-B) and basal-to-apical (B-A) permeability of CI-994, W2A-28, W2A-42 and W2A-43 in the presence of cyclosporine A.

| Compound | P<sub>app</sub> (A-B), 10<sup>-6</sup> cm/s |  |  | P<sub>app</sub> (B-A), 10<sup>-6</sup> cm/s |  |  | Efflux ratio* |
| --- | --- | --- | --- | --- | --- | --- | --- |
|  | Data1 | Data2 | Mean | Data1 | Data2 | Mean |  |
| Quinidine | 18.49 | 17.03 | **17.76** | 12.31 | 15.61 | **13.96** | **0.79** |
| Digoxin | 0.27 | 0.31 | **0.29** | 1.67 | 1.80 | **1.74** | **6.04** |
| CI-994 | 9.92 | 9.91 | **9.91** | 10.89 | 10.54 | **10.71** | **1.08** |
| W2A-28 | 20.82 | 20.51 | **20.67** | 15.29 | 18.90 | **17.10** | **0.83** |
| W2A-42 | 22.90 | 24.12 | **23.51** | 19.97 | 16.30 | **18.13** | **0.77** |
| W2A-43 | 5.20 | 7.22 | **6.21** | 8.39 | 7.95 | **8.17** | **1.32** |

\* Efflux ratio is expressed as the quotient of P<sub>app</sub>(BA) to P<sub>app</sub>(AB).

### W2A-28 induces histone acetylation and activates Wnt/β-catenin signaling in AD patient-specific iPSC-derived cerebral organoids

Cerebral organoids derived from AD patient-specific iPSCs can represent multifactorial phenotypes of AD and are excellent disease models for AD research and drug development (41-43). Therefore, cerebral organoids derived from two AD patient-specific iPSC lines were used for the current studies. The iPSC line MC0020 was derived from a female patient with late-onset AD carrying *APOE* ε4/ε4 genotype, and the iPSC line MC0127 was derived from a female patient with familial AD carrying *APOE* ε3/ε4 genotype and *PSEN1* M146V mutation. The validation of the iPSC line MC0020 on its pluripotency and karyotype has been reported (32). The pluripotency of the iPSC line MC0127 was confirmed by the expression of pluripotency markers TRA-1-60 and Nanog (***Fig. S1A***), and its differentiation capacity was verified by the ability to differentiate into endodermal (Sox17^+^), mesodermal (Brachyury^+^), and ectodermal (Nestin^+^) cells (***Fig. S1B***). Karyotyping test confirmed that the number and appearance of chromosomes were normal in iPSC line MC0127 (***Fig. S2***).

To determine the effects of W2A-28 on class I HDAC inhibition and Wnt/β-catenin signaling activation, the levels of histone H3 acetylation and β-catenin were examined in cerebral organoids derived from AD patient-specific iPSC lines MC0020 and MC0127 at 3 months of age. It was found that W2A-28 at 1 and 2 μM greatly increased the levels of histone H3 acetylation in AD patient-specific iPSC-derived cerebral organoids carrying *APOE* ε4/ε4 or *APOE* ε3/ε4 with *PSEN1* M146V mutation (***Fig. 5A & 5B***). Moreover, W2A-28 significantly increased total cellular β-catenin levels in AD patient-specific iPSC-derived cerebral organoids (***Fig. 5C & 5D***). *NEUROD1* is a specific target gene of Wnt/β-catenin signaling in the central nervous system (CNS) (44). Western blotting studies revealed that the level of NeuroD1 protein was significantly increased upon W2A-28 treatment in AD patient-specific iPSC-derived cerebral organoids (***Fig. 5C & 5D***). Together, these findings indicate that W2A-28 increases histone acetylation and activates Wnt/β-catenin signaling in AD patient-specific iPSC-derived cerebral organoids.

**Fig. 5.**
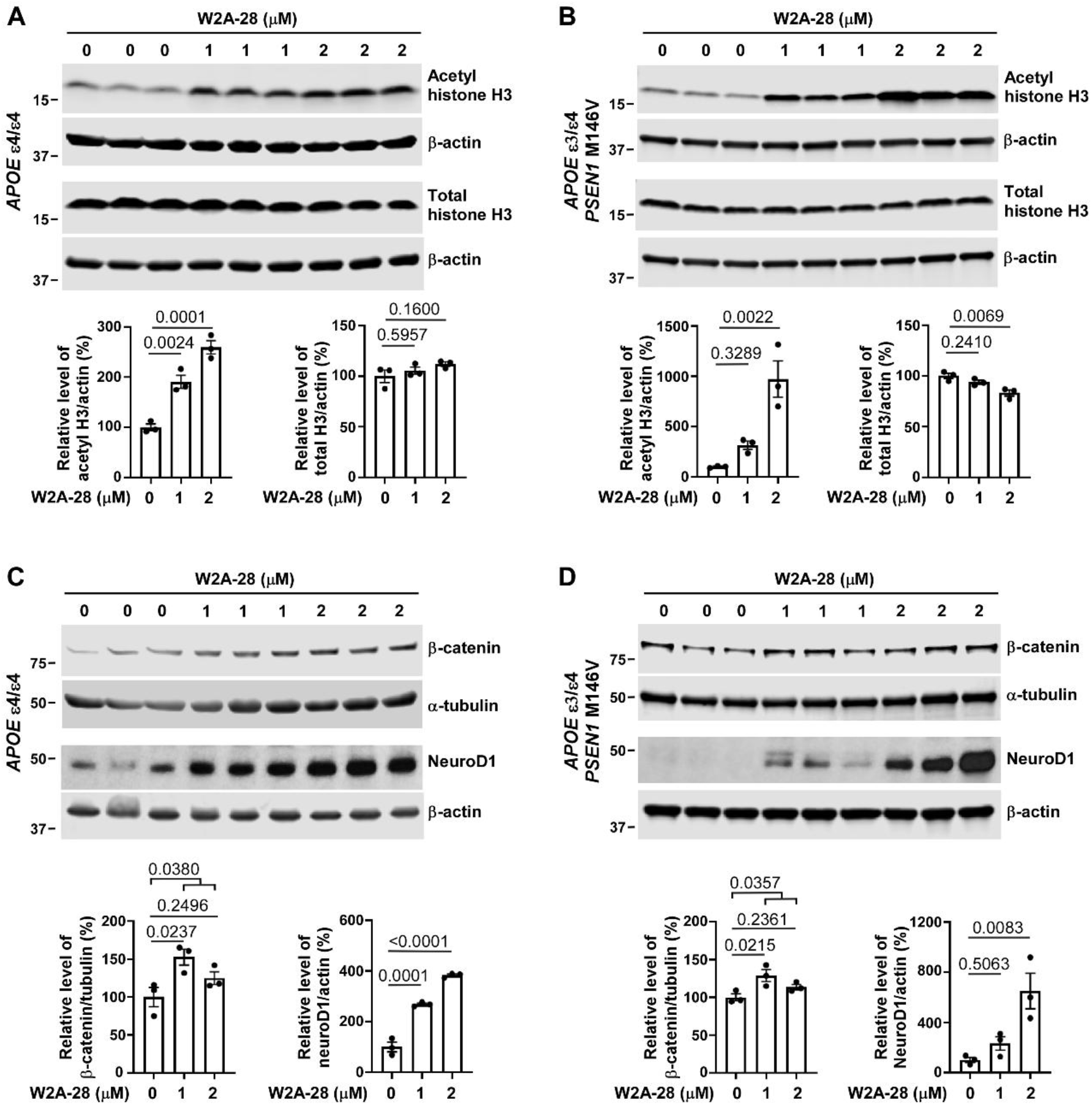
W2A-28 enhances histone H3 acetylation and activates Wnt/β-catenin signaling in AD patient-specific iPSC-derived cerebral organoids. Human cerebral organoids carrying *APOE* ε4/ε4 (A, B) or *APOE* ε3/ε4 with *PSEN1* M146V mutation (C, D) at 3 months of age cultured in 6-well plates (3 organoids per well) were treated in triplicate with CI-994 at indicated concentrations for 72 h. The levels of acetyl histone H3K9, total histone H3, total β-catenin, β-actin, NeuroD1 and α-tubulin were examined by Western blotting and quantified. Data represent mean ± SEM. *P* values were calculated using one-way ANOVA with Dunnett’s multiple comparison test or two-tailed, unpaired t test.

### W2A-28 significantly inhibits tau phosphorylation in AD patient-specific iPSC-derived cerebral organoids

Hyperphosphorylation of the microtubule-associated protein tau in neurons plays a key role in neurodegenerative tauopathies in AD (45, 46). To determine the therapeutic effects of W2A-28 in AD, tau phosphorylation was examined in AD patient-specific iPSC-derived cerebral organoids at 3 months of age. Western blotting studies revealed that W2A-28 at 1 and 2 μM significantly reduced tau phosphorylation, which was detected by two well-known tau phosphorylation antibodies AT8 and AT180, in AD patient-specific cerebral organoids carrying *APOE* ε4/ε4 (***Fig. 6A-E***) or *APOE* ε3/ε4 with *PSEN1* M146V mutation (***Fig. 6F-J***). Together, these findings indicate that W2A-28 treatment results in a significant inhibitory effect on tau phosphorylation in AD patient-specific iPSC-derived cerebral organoids.

**Fig. 6.**
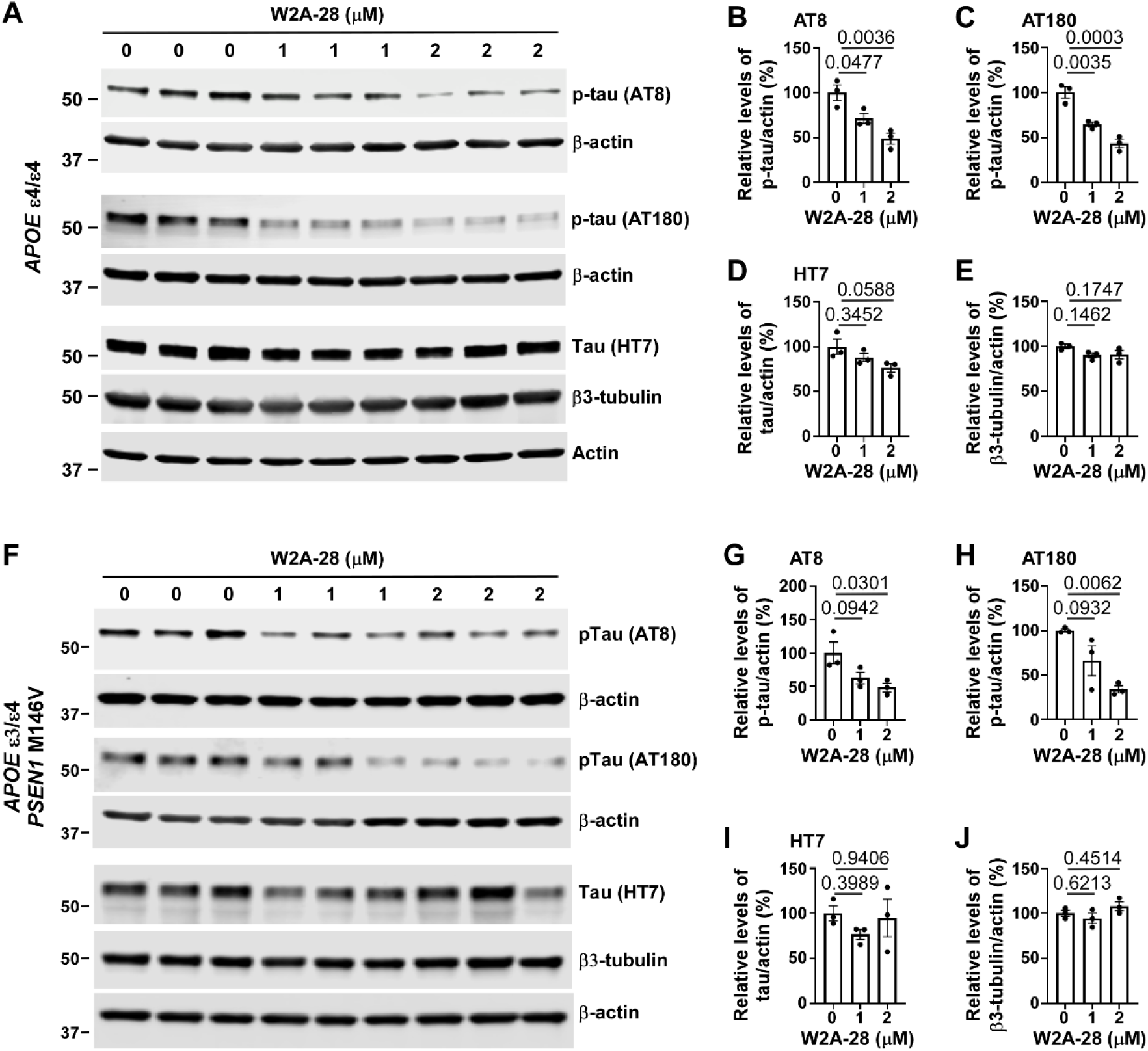
W2A-28 suppresses tau phosphorylation in AD patient-specific iPSC-derived cerebral organoids. Related to Fig. 4, human cerebral organoids carrying *APOE* ε4/ε4 (A-E) or *APOE* ε3/ε4 with *PSEN1* M146V mutation (F-J) at 3 months of age cultured in 6-well plates (3 organoids per well) were treated in triplicate with W2A-28 at indicated concentrations for 72 h. Western blotting analyzed the levels of phosphor-tau detected by AT8 or AT180 antibodies, total cellular tau detected by HT7 antibody and neuronal marker detected by β3-tubulin antibody, and normalized with β-actin levels. All the values are mean ± SEM. *P* values were calculated using one-way ANOVA with Dunnett’s multiple comparison test.

## Discussion

CI-994 is an orally bioavailable specific class I HDAC inhibitor and a brain penetrant (19, 20), and clinical trials for several types of cancer have revealed that CI-994 is well tolerated in humans (16-18). CI-994 is a dual modulator of class I HACs and Wnt/β-catenin signaling, works as a molecular memory aid and alleviates cognitive impairment and amyloid and tau pathologies in AD mouse models and AD patient-specific iPSC-derived cerebral organoids (8, 14, 20-24). Therefore, CI-994 is an ideal scaffold for further development of AD drug candidates. Here we discover W2A-28, a derivative of CI-994, as a potential better candidate than CI-994 for AD treatment. W2A-28 functions as a dual modulator for class I HDAC inhibition and Wnt/β-catenin signaling activation and significantly suppresses tau phosphorylation in AD patient-specific iPSC-derived cerebral organoids carrying *APOE* ε4/ε4 or *APOE* ε3/ε4 with *PSEN1* M146V mutation.

Class I HDACs, which comprise HDAC1, 2, 3, 8, are epigenetic master regulators of chromatin function. Inhibitors of class I HDACs can significantly improve cognitive function in various models of neurological diseases (8, 9, 22). While HDAC2 and HDAC3 are neurotoxic (47, 48), HDAC1 displays both neurotoxic and neuroprotective roles (49). Mounting evidence indicates that HDAC3 is an attractive therapeutic target for disease-modifying therapy in AD (48). CI-994 specifically inhibits HDAC1, 2 and 3 activities with IC_50_ values of 0.62 μM, 0.59 μM and 0.47 μM, respectively (14). Interestingly, W2A-28 inhibits HDAC1, 2 and 3 activities with IC_50_ values of 0.51 μM, 0.68 μM and 0.22 μM, respectively, and has no inhibitory effects on HDAC8 and other HDACs, indicating that W2A-28 has some selectivity for HDAC3 inhibition (2.3 – 3.0-fold selectivity over HDAC1 and HDAC2). Therefore, W2A-28 could be a better class I HDAC inhibitor than CI-994 for AD therapy. It is worth noting that CI-994 derivative W2A-42 specifically inhibits HDAC1, 2 and 3 activities with IC_50_ values of 1.51 μM, 1.53 μM and 0.33 μM, respectively, indicating that W2A-42 is a specific HDAC3 inhibitor with about 4.6-fold selectivity over HDAC1 and HDAC2. Further studies are required to determine its therapeutic effects in AD.

Wnt/β-catenin signaling is greatly suppressed in AD brain via multiple pathogenic mechanisms, which contributes to AD pathogenesis (15). Notably, the expression of Wnt co-receptor LRP6 is downregulated in AD brains (50) and two LRP6 SNPs and an alternative splice variant, which display impaired the activity of Wnt/β-catenin signaling, are associated with synapse loss and increased risk of developing AD (51-53). Indeed, neuronal LRP6 deficiency leads to synaptic dysfunction and exacerbates amyloid pathology in AD (50). In the present study, we have found that W2A-28 is more potent than CI-994 in activating Wnt/β-catenin signaling with an EC_50_ value of 1.61 μM in Wnt-3A-expressing HEK293 cells. Mechanistically, W2A-28 and CI-994 as well activate Wnt/β-catenin signaling by LRP6 stabilization. In addition, there is a potential therapeutic-positive feedback loop between class I HDAC inhibition and Wnt/β-catenin signaling activation (14). Our findings strengthen the notion that restoring LRP6-mediated Wnt/β-catenin signaling is a potential effective strategy for AD.

As a specific target of Wnt/β-catenin signaling, the basic helix-loop-helix transcription factor *NEUROD1* is upregulated upon activation of Wnt/β-catenin signaling in CNS (44). Interestingly, NeuroD1-based gene therapy demonstrates therapeutic potential in mouse models of AD and other neurological disorders (54-57). NeuroD1 treatment greatly enhances neurogenesis, attenuates reactive astrocytes-mediated neuroinflammation and improves motor and cognitive functions *in vivo* (54-57). In our recent study, we have shown that CI-994 remarkedly enhances *NEUROD1* expression in AD patient-specific iPSC-derived cerebral organoids (14). In the present study, we have demonstrated that W2A-28 dramatically increased NeuroD1 protein levels in AD patient-specific iPSC-derived cerebral organoids. Therefore, induction of *NEUROD1* expressing could be an important mechanism associated with CI-994/W2A-28-mediated AD therapy.

Membrane permeability and Pgp ae restrictive factors for CNS drugs to cross BBB. For CNS penetration, a drug should ideally have an *in vitro* passive permeability >15 ×10^−6^ cm/s and not be a good Pgp substrate (efflux ratio <2.5) (58). It has been demonstrated that CI-994 can cross BBB [16]. In the present study, we have demonstrated that both CI-994 and W2A-28 display excellent metabolic stability in mouse and human liver microsomes. While both CI-994 and W2A-28 are not Pgp substrates, the level of W2A-28 permeability coefficient is more than 2-fold of that of CI-994 permeability coefficient in MDR1-MDCKII cells, suggesting that W2A-28 is a potential brain penetrant. Further studies are required to study its PK profile and brain penetration *in vivo*, and determine its therapeutic effects in mouse models of AD.

## Authors’ contributions

Y.L. developed the research concept and designed the experiments. T.K. and G.B. contributed to scientific discussions. T.R.C., Y.L. designed CI-994 derivatives and coordinated chemical synthesis. T.R.C. coordinated metabolic stability and permeability studies. W.L., E.L., S.J., Y.L. conducted biological experiments and/or data analyses. The manuscript was drafted by Y.L., and edited by G.B. and T.R.C. All authors read and approved the manuscript.

## Acknowledgment

We are grateful to Dr. Steve Younkin for helping with the collection of human skin biopsies for generating the iPSC line MC0020 and MC0127. This work was supported in part by the grants R21AG065653 and R01AG078615 from the National Institutes of Health (to Y.L.) and the grant 24A07 from the Florida Department of Health Ed and Ethel Moore Alzheimer’s Disease Research Program (to Y.L.).

## Declarations of conflict of interest

G.B. consults for SciNeuro Pharmaceuticals. T.R.C. is a current employee of Digital Ether Computing. All other authors declare no competing interests.

