## Supplemental Figures for "Discovery of a CI-994 derivative as a dual modulator of class I HDACs and Wnt/β-catenin signaling for Alzheimer’s disease therapy"

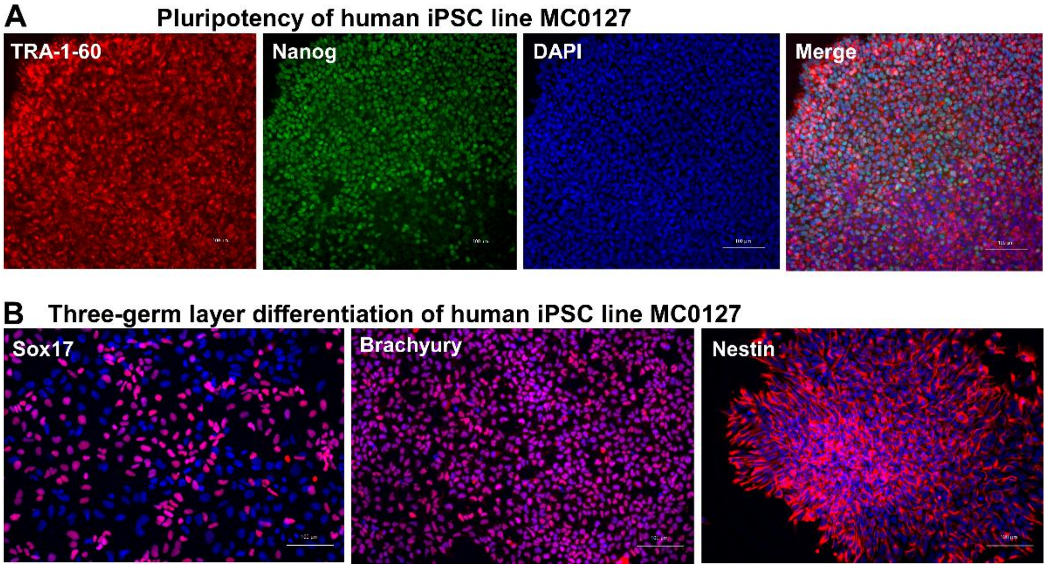


**Fig. S1. The pluripotency of the iPSC line MC0127.** (A) Immunostaining of pluripotency markers (TRA-1-60 and Nanog) in iPSC line MC0127 carrying *APOE* ε3/ε4 and *PSEN1* M146V. (B) *In vitro* differentiation of iPSC line MC0071 into cells of all three germ layers. Cells were for Sox17 (endoderm), Brachyury (mesoderm), Nestin (ectoderm), and DAPI (nucleus). Scale bar:100 µm.

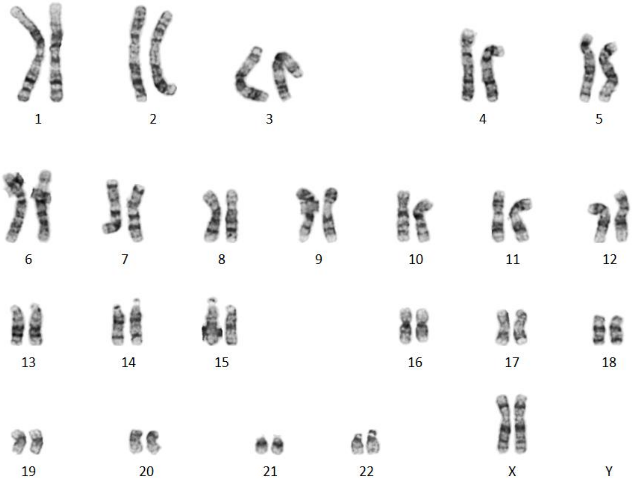


**Fig. S2. Karyotyping analysis of the iPSC line MC0127.**
